# Cortical Activity Associated With Acute Development of Phantom Auditory Percepts via Unilateral Deprivation

**DOI:** 10.1101/2023.06.13.544737

**Authors:** Gregory Ginnan, Stefan Rampp, Roland Schaette, Michael Buchfelder, Nadia Müller-Voggel

## Abstract

Subjective tinnitus describes the experience of hearing phantom sounds (e.g., tones, buzzing, noise). While the majority of those who report having experienced phantom sounds claim that these percepts have not lasted but are transient, some experience this chronically, with others even describing their tinnitus as severe enough to negatively impact their well-being and daily lives. Currently, no permanent solution has been discovered for preventing or curing tinnitus. This is due to several factors, including an insufficient understanding of the mechanisms at play that give rise to such auditory sensations, as well as a lack of research investigating the corresponding changes in neural activity associated with the onset and development of tinnitus. Taking advantage of the high spatial and temporal resolution of magnetoencephalography (MEG), we measured cortical activity associated with the development of acute tinnitus-like percepts induced via unilateral auditory deprivation. Over the course of four days, participants continuously wore a silicone earplug in one ear, which led to the experience of phantom sounds in 15 of 16 participants. Frequency analysis of source-localized continuous MEG data revealed a significant increase of gamma power in primary auditory cortices (A1) during the tinnitus condition (p=0.02), which most likely reflects the neuronal processing correlated with tinnitus perception.

## 1. INTRODUCTION

Although the ubiquity of subjective tinnitus in the general population has become increasingly clear over recent years (McCormack et al., 2016), future tinnitus prevalence is likely to rise, given increased rates of urbanization and personalized headphone use, both of which contribute to louder, more intense acoustic environments for individuals, a risk factor for the development of tinnitus. Despite growing interest in and attention towards tinnitus within the neuroscientific community, many of the underlying neural mechanisms remain unresolved. Many published neuroimaging studies have focused on populations whose tinnitus can be considered “chronic” (i.e., tinnitus duration > 1 month) (Weisz et al., 2005, 2007; Sedley et al., 2012; Seydal et al., 2013; Adjamian et al., 2014), while relatively few have used neuroimaging techniques such as magnetoencephalography (MEG), electroencephalography (EEG), or functional magnetic resonance imaging (fMRI) to capture cortical activity associated with the development of acute tinnitus in humans. This may be in part due to the difficulty in finding research participants whose tinnitus has only lasted for a few days or weeks. Alternatively, this tendency in tinnitus neuroimaging studies to report activity mostly from chronic tinnitus sufferers may reflect the urgency associated with finding a solution for that population in particular. A third possible explanation for the lack of cognitive neuroscience papers investigating acute tinnitus development is most likely related to the scarcity of safe and reliable methods to induce a temporary phantom auditory sensation.

There are methods known to induce tinnitus in other mammals, but are neither approved nor safe for application in humans. One such technique is the use of salicylate to initiate a temporary hair-cell dysfunction (Eggermont 2012; Lanaia et al., 2021). Another method is to expose animals to acoustic noise trauma, with sound stimuli reaching and maintaining decibel levels sufficient to disrupt cochlear function via damage-induced hearing loss (Tziridis et al., 2015). These methods are for good reason prohibited when it comes to human study, therefore leaving only a few limited, yet interesting approaches at researchers’ disposal. For example, one MEG study monitored oscillatory activity from participants before and immediately after they attended their respective rock band rehearsals, in an attempt to see what trace a (voluntary) sustained exposure to elevated decibel levels may leave in cortical oscillatory patterns (Ortmann et al., 2011).

Deprivation is a method researchers use to induce temporary plastic changes in the human auditory system to simulate hearing loss. This may be achieved via exposure to silence in a soundproof booth or anechoic chamber, or by employing an earplug or physical blockage of some kind to reduce incoming noise levels. Most deprivation studies which use earplugs, both unilateral and bilateral, focus on other changes in non-tinnitus sensory experiences associated with reduced cochlear input, such as hyperacusis and hearing loss (Decker and Howe, 1981; Formby et al., 2003, 2007; Munro and Merrett, 2013; Maslin et al., 2013; Brotherton et al., 2016, 2017). None of these studies reported detrimental side effects linked to deprivation in their participants. Unilateral deprivation has also been proven capable of eliciting tinnitus-like phantom auditory percepts within only a few days, and in the existing literature no serious side effects related to the development of these auditory sensations have been reported, although that list is admittedly small (Schaette et al., 2012; Brotherton et al. 2019). In their investigation, Schaette et al. used silicone earplugs to simulate unilateral hearing loss in 18 participants with normal hearing. During the course of 7 days of continuous earplug use, 14 participants (78%) experienced phantom auditory sensations. Indeed, sound attenuation via external earplug does seem a promising approach with which to temporarily mimic changes associated with hearing loss, and was the main method employed in this study.

### 1.1 Tinnitus-Related Changes of Auditory and Non-auditory Systems

Tinnitus has been associated with plastic changes at different points along the auditory pathway, most usually observed as some form of hyperactivity, such as an increase in neural synchrony (Eggermont and Tass, 2015) or elevated spontaneous firing rate levels (Kiang et al., 1970; Brozoski et al., 2002). Neural synchrony occurs when individual neurons rhythmically fire at the same moment, leading to local field potential synchronization which becomes detectable by EEG or MEG as a neural oscillation. Previous neuroimaging studies have investigated the oscillatory activity associated with chronic tinnitus (Weisz et al., 2005, 2007; Sedley et al., 2012; Demopoulos et al., 2020). One EEG study demonstrated that percept loudness was correlated with heightened levels of contralateral primary auditory cortex (A1) gamma activity in chronic unilateral tinnitus patients (Van der Loo et al., 2009), while another found lateralized gamma power “hot spots” on unilateral tinnitus patients’ temporal lobes compared to healthy controls (Ashton et al., 2007). Using MEG, Weisz et al. revealed not only significantly increased auditory cortical gamma activity in chronic tinnitus patients, but that this gamma activity was correlated with the onset of delta waves (Weisz et al., 2005). This same chronic tinnitus group also displayed significantly decreased alpha activity when compared with healthy participants. Adjamian et al. revealed elevated A1 delta-band power in tinnitus sufferers’ MEG resting state data, but only in those with both tinnitus and hearing loss (Adjamian et al., 2012; Shore et al., 2016), while another MEG study examining tinnitus sufferers’ A1 also found increased delta and gamma oscillations, alongside reduced alpha oscillations (Wienbruch et al., 2006). In patients with chronic tinnitus who underwent six months of tinnitus retraining therapy (TRT), gamma power was significantly reduced in the left A1 and A2, correlating with reported reduced stress and perception (Lee et al., 2019). In one study, changes in EEG power spectra were monitored as participants who could control or “turn on” their tinnitus via various strategies (e.g., tooth/jaw clenching) (Zhang et al., 2021). Increased delta, theta, and gamma power were observed in the tinnitus-on condition, with alpha power showing wide variability between participants. A possible explanation for the relationship of these altered auditory cortical patterns with the experience of tinnitus is that alpha, often associated with inhibition of task-irrelevant areas (Knyazev, 2007), if decreased, could result in disinhibition of cortical areas responsible for encoding auditory stimuli. This disinhibition would then allow cells, which would otherwise be silenced, to fire and eventually synchronize in the form of gamma oscillations, often giving rise to a conscious auditory percept. Although tinnitus is primarily thought of as a pathological state of the auditory system, a number of non-auditory brain structures have been shown to exhibit altered spontaneous oscillatory activity in tinnitus sufferers (Vanneste & De Ridder, 2012). Given that tinnitus in general, and chronic tinnitus in particular, may often lead to feelings of distress or frustration, it is perhaps not surprising that brain regions responsible for emotional regulation, namely limbic and prefrontal areas, are among the usual cortical suspects implicated in tinnitus distress networks (Lockwood et al., 1998; Rauschecker et al., 2010; Milner et al., 2020; Kanzaki et al., 2021). Additionally, cortical regions responsible for attention, memory, and somatosensory processing are also thought to be involved in chronic tinnitus networks. Increased delta oscillations correlated with phantom sound perception in chronic tinnitus sufferers have been observed not just in A1, but also across temporal, parietal, sensorimotor, and limbic cortices (Sedley et al., 2015). This has led to speculation that perhaps altered slow-wave activity (SWA) in chronic tinnitus is related to increased communication between auditory and non-auditory areas, which could lead to a reinforcement or entrenchment of tinnitus percepts (Shore et al., 2016). The role of neural oscillations seems to be central to tinnitus network activity, as studies have shown significantly altered alpha-band long-range coupling between parietal, prefrontal, and auditory cortical structures (Schlee et al., 2009a). The subjective intensity of tinnitus distress has also been linked to changes in cortico-cortical coupling as measured by information flow directionality, particularly above 30 Hz (Schlee et al., 2009b). Additionally, functional connectivity has also been shown to be increased between limbic and auditory regions in tinnitus patients versus controls (Chen et al., 2017). Therefore, although tinnitus may be perceived as a simple sound, it is becoming clear that these phantom auditory percepts are the conscious representation of a much more complicated interplay of cortical and subcortical structures, both auditory and non-auditory. These local and global networks seem to rely on oscillatory dynamics which, if disrupted, may influence the tinnitus percept or its associated emotional component (Riha et al., 2020).

### 1.2 Neural Oscillations in Acute Tinnitus

As already mentioned, the bulk of neuroscientific literature dealing with tinnitus investigates those with chronic tinnitus rather than acute. However, a few studies have focused on people whose tinnitus’ onset is relatively recent (less than one month). One experimental design measured cortical oscillatory activity in rock musicians before and after band rehearsal, during which they were exposed to an average loudness level of 97.3 dB SPL, and who reported experiencing a transient tinnitus following rehearsal (Ortmann et al., 2010). Compared to controls, the musicians’ MEG data revealed significantly increased gamma power (55-85 Hz) lateralized to the right auditory cortex. However, the authors refrained from explicitly claiming a causal link between increased gamma activation and the tinnitus percept itself. A recent EEG study found increased gamma-band activity (55-100 Hz) localized to the middle frontal gyrus and the parietal gyrus in acute tinnitus sufferers compared to chronic tinnitus sufferers (Lan et al., 2020). When compared with healthy controls, the acute tinnitus group demonstrated decreased activity across frequency bands in the superior frontal cortex, whereas the chronic tinnitus group showed a reduction in the same region, but only in the beta (21.5-30 Hz) and gamma (55-100 Hz) bands. Interestingly, no changes in neural oscillations were found in auditory cortices in any frequency band between acute and chronic tinnitus groups, nor when compared with healthy controls. Lan et al. also showed a significant increase in functional connectivity between auditory and non-auditory areas in the chronic tinnitus group compared with the acute tinnitus group, suggesting that the process of phantom auditory percept chronification involves increased crosstalk among several brain regions.

Using MEG, we have sufficient temporal and cortical spatial resolution to explore these oscillatory phenomena and how they might relate to the conscious auditory experiences with no external sound sources. By earplugging one ear for four days, we were able to monitor changes in neural oscillations associated with artificial hearing loss, plasticity of the auditory pathway, and, for all but one of our participants, the experience of tinnitus-like percepts on the plugged side. This study focused on oscillatory activity originating from primary and secondary auditory cortices. When participants were actively experiencing phantom auditory percepts, we expected to find oscillatory dynamics similar to those found in chronic tinnitus sufferers, namely an increase in both delta (1-4 Hz) and gamma (30-60 Hz) power, accompanied by a concomitant decrease in alpha (8-14 Hz). Future analyses will focus on connectivity between auditory and non-auditory areas to determine the influence of other cortico-cortical modulations which might affect auditory cortical activity.

## 2. METHODS

We recruited a total of 17 right-handed participants who were required to have normal hearing, no inter-ear asymmetry greater than 5 dB, as well as no history of tinnitus or hyperacusis (10 male, 7 female, mean age = 26.5 ± 2.6). Of the 17 participants, 12 completed the entire experiment (whose results will be discussed here), four partially completed, and one participant failed to wear the earplug for the full four days. Therefore, our final dataset is comprised of 12 participants (8 male, 4 female, mean age = 26.7 ± 2.6), while the ear being plugged was divided between left (5) and right (7). Patients also filled out the ICD-10-Symptom-Rating (ISR) psychological questionnaire (Tritt et al., 2008).

All participants provided their written informed consent. The study was positively reviewed by the Institutional Review Board (IRB) (392_18 B) of the medical faculty of the Friedrich-Alexander-University Erlangen-Nuremberg, Germany.

### 2.1 Audiometry

Participants’ hearing thresholds were measured via pure-tone audiometry with a SentiFlex Audiometer (PATH Medical, Germering, BY, Germany). Audiometry was carried out in line with ISO 8253-1 procedures. During the hearing tests, participants sat in a 250 Series Mini Sound Shelter (Industrial Acoustics Company GmbH, Niederkruechten, Germany), fitted with HDA-280 headphones (PATH Medical, Germering, BY, Germany). Each frequency was presented with an initial decibel level of -20 dB HL. Given a trigger from the participant, intensity was subsequently decreased by steps of 10 dB until no trigger was delivered (i.e., the participant did not hear a tone). Intensity was then increased in 5-dB intervals until the tone was again detected by the participant, and the process repeated until the hearing threshold for a given frequency was determined.

Audiometry was performed a total of four times per participant, both before and after each of the two MEG measurements. The initial audiological test also served as a screening exam in order to ensure participants’ hearing in both ears fell within what is considered a healthy range (≤ 20 dB HL from 0.25-8 kHz). Participants were only included if their inter-ear hearing threshold asymmetry was less than 5 db. Neither average hearing thresholds nor inter-ear differences were significantly altered during the course of the experiment (Table 1).

**Table 1.** Characteristics of subjective phantom percepts.

| Participant Number | Plugsides | Tinnitus during plug period | Tinnitus on day 4 | Tinnitus description | Estimated frequency (Hz) |
| --- | --- | --- | --- | --- | --- |
| 1 | R | Y | Y | Tone | 4750 |
| 5 | L | Y | Y | Noise, beeps, whistling | n/a |
| 6 | R | Y | N | Tone | n/a |
| 7 | L | Y | Y | High tone, low buzz | 7200 |
| 8 | R | Y | Y | Tone | 4200 |
| 9 | L | Y | Y | Noise | n/a |
| 10 | R | Y | Y | Tone | 6000 |
| 11 | R | Y | Y | Tone | 6500 |
| 15 | R | Y | Y | Tone (day), Noise (night) | 5500 |
| 16 | L | Y | Y | Tone, buzzing | 2000 |
| 17 | R | N | N | n/a | n/a |
| 18 | L | Y | Y | Tone, beeping | 2100 |

We also used the audiometer to measure the effectiveness of the earplugs in attenuating incoming noise throughout the experiment. At the end of the first MEG measurement, participants inserted the plug for the first time, and by conducting another audiometric test as described above, we verified that detection thresholds were decreased to decibel levels that correspond with those of mild or moderate hearing loss (25-40 dB HL and 40-55 dB HL, respectively) (Table 2). The average attenuation corresponded to vendor specifications (Mack’s Earplugs, McKeon Products, Inc., MI, USA) (Table 3).

Audiometry with the earplug still inserted was also performed when the participants returned after 4 days of consecutive plug use. Thus, we ensured that the plug was still attenuating outside noise prior to the second MEG measurement. The final audiometric reading to verify noise attenuation was taken just before the final (i.e., recovery) MEG measurement.

### 2.2 Magnetoencephalography (MEG)

Participants’ brain activity was recorded during two magnetoencephalographic measurements using a Magnes 3600 WH MEG system (248 magnetometers, 4D Neuroimaging, San Diego, CA, USA). Data were recorded at a sampling frequency of 678.2 Hz, with an online high-pass filter of 1.0 Hz and a low-pass filter of 200 Hz. Ambient noise was corrected for using a calibrated linear weighting of 23 reference sensors (manufacturer’s algorithm, 4D Neuroimaging, San Diego, CA, USA). Prior to the beginning of each scan, participants’ headshapes were digitized with a 3D-digital pen (Polhemus, Colchester, Vermont, USA). Typically, around 300-500 points were collected with the pen, ranging along the brow, scalp, and nose. Next, the digital pen was used to mark 5 anatomical points (nasion, LPA, RPA, Cz, inion), as well as the position of 5 fixed head coils which so as to monitor head movement. The marking of the anatomical points and head coils were used in combination with the digitized headshape for later co-registration steps necessary for source reconstruction.

### 2.3 Experimental Design

In order to record and compare the MEG activity of participants during and after the development of acute tinnitus-like percepts, participants underwent two MEG measurements (hereafter referred to as Day 4 and Recovery measurements). During participants’ first visit to our lab, they were fitted with the earplug (randomized between left (5) and right (7) for the first time, undertook a final hearing test with the plug inserted to ensure noise attenuation, and then sent home until their return to the lab four days later.

These two MEG measurements were part of a larger study including more resting state measurements while participants listened to auditory tone simulation. To avoid influence on the measurements described in this study, these additional MEG data were recorded at the end of each experimental session.

#### 2.3.1 Day 4

After having worn the earplugs consecutively for four days, participants returned to the lab, with the earplug still inserted. As described above (*Audiometry*), they underwent an audiometric exam upon arrival to ensure continued attenuation by the plug. Next, participants were asked to match their phantom auditory percept to similar tones played through a headphone to the unplugged ear using the open-source digital audio editor Audacity (https://audacityteam.org/). Pure tones were periodically presented to the unplugged ear, starting at an initial frequency of 2 kHz at 10 dB SPL. According to the participants’ judgments, the frequency was either increased or decreased by 1 kHz in the direction of their subjective tinnitus frequency. Subsequent changes to frequency were done increasingly smaller increments, so as to adjust the stimulated ear’s pure tone to match the plugged ear’s perceived tinnitus pitch as closely as possible. Loudness matching was not performed, with the presented pure tones fixed at -10 dB SPL. After these steps and with the plug still inserted, participants were placed inside the MEG and underwent two five-minute resting state recordings, the first while wearing the earplug, and the second without. This first resting state measurement was critical to the experimental setup, as this served as our “tinnitus” condition; during this measurement, we expected most participants to be experiencing unilateral tinnitus-like sensations on the ear ipsilateral to the plugged side (indeed, 10 out of 12 participants reported tinnitus-like percepts during this particular measurement).

#### 2.3.2 Recovery

The experimental protocol of the second MEG measurement, termed the recovery measurement (given that at this point (2-4 weeks post-plug removal), all participants had experienced a disappearance of the tinnitus-like phantom sounds), was the same as Day 4 (five-minute plugged resting state, only this time without tinnitus-like percept in all participants). This measurement was necessary to compare resting states while the earplug was being worn, so as to account for any differences attributable to hearing loss.

One participant (P13) who completed the measurements on Day 1 and Day 4 did not take part in the recovery measurement, as their tinnitus did not disappear as expected, and therefore has been excluded from all analyses. More information is provided below in the section entitled *Role of Attention in Tinnitus*.

### 2.4 Earplug Care and Maintenance

After discussions with the researchers who previously induced tinnitus via unilateral deprivation (Schaette et al., 2012), as well as taking part in our own pilot study during which the plug was worn for one week, we determined that four days of earplug use would be sufficient to elicit a tinnitus-like percept. Once fitted with the earplug at the lab, participants were instructed to wear the earplug in the same ear continuously for four days, for ∼23 hours per day. They were only permitted to remove the plug for daily hygienic purposes, particularly during bathing or showering. Each participant was given 5 earplugs so they could swap out a clean plug for each day’s previous one. Participants were also given a series of tips that would ease the annoyance that could come with wearing a plug in only one ear, such as the amplified volume (due to bone conduction) of one’s own voice, breath, eating, and drinking, as well as safety tips to ensure / remind participants that they must remain vigilant and aware while biking or walking around outside, as half of their auditory input would be significantly lessened.

### 2.5 Daily Journal

Participant were also given a daily journal in which they were encouraged to record both physiological (auditory) and emotional observations throughout the experiment. We wanted participants to feel comfortable enough to share with us what went through their minds during this time of experiencing tinnitus-like percepts, as emotional distress has been shown to be associated with increased tinnitus severity (this was also communicated to the participants, per the study’s ethics proposal) (Malouff et al., 2011). On the auditory side, participants were encouraged to note if and when phantom auditory sounds began to appear. If they were able, they were also encouraged to make note of other sonic characteristics of the developing tinnitus-like percept, such as pitch (frequency), loudness (intensity), or timbre (sonic color or texture, giving rise to individual or distinct sounds (e.g., difference between saxophone and trumpet)). Participants were also asked to record the time when these emotional or auditory sensations arose, so as to give us a better idea if any patterns throughout the day and night could be influencing the participants’ experience and/or perception. The degree to which the participants engaged with and wrote entries in the daily journal varied widely among the participants, with some writing a couple of words each day, while others found the one sheet of paper provided insufficient, returning five pages of their own observations.

## 3. Data Analysis

### 3.1 Audiometric Data

#### 3.1.1 Hearing Thresholds

Participants’ hearing thresholds derived from pure-tone audiometry were statistically compared using paired-sample Student’s t-tests to ensure a) distinct audiometric profiles of the plugged and unplugged ears, b) unaltered earplug attenuation of the plugged ear over the course of the experiment, and c) no difference in hearing sensitivity of either ear upon completion of the experiment (i.e., recovery to baseline thresholds).

#### 3.1.2 Hearing Loss

We tested whether the frequencies of participants’ subjective tinnitus-like percepts were modulated by overall hearing loss. Hearing loss was determined by first averaging the hearing thresholds of the plugged ear as measured at the end of Day 1 and the beginning of Day 4. Next, the areas under the audiometric curves (AUCs) for the plugged ear were computed, both before and during earplug use. The AUC of the averaged plugged audiogram was then subtracted from the initial audiogram of the same ear, giving an overall hearing loss value in decibels per octave for each participant. We tested for linear correlation between hearing loss and subjective percept frequency using Pearson’s correlation.

### 3.2 MEG Signal Acquisition and Analysis

The majority of the following analyses, unless noted otherwise, were performed using the MNE-Python toolbox (Version 3.7.10, Gramfort et al., 2013). Raw MEG data were downsampled from 687.2 Hz to 200 Hz and notch-filtered at 16 2/3 Hz, 50 Hz, and 100 Hz (passing trains, 50-Hz electrical line noise, electrical line noise harmonic, respectively). Next, the downsampled and filtered data were separated into 2-second epochs. Epochs containing eye blinks and movement artifacts were dropped, with the remaining good epochs run through an independent component analysis (ICA) using the Picard method (preconditioned ICA for real data) to identify and remove cardiac signatures and other possible contaminations of the data (e.g., external noise). Next, participants’ headshapes were co-registered to a common source space (called “fsaverage” brain), provided by the FreeSurfer software suite, which is documented and freely available for download online (http://surfer.nmr.mgh.harvard.edu/). This step yielded a bilateral hemisphere (4096 vertices per hemisphere) surface-based source space per participant, using the default spacing of a recursively subdivided octahedron. A boundary element model per participant was computed using MNE-Python’s linear collocation approach. This was then used to compute leadfields per participant and condition. Leadfields per participant were averaged and used as a common leadfield only if head position had moved less than 5 mm between conditions or experimental days.

Cross-spectral density (CSD) matrices were constructed by convolving time series data with complex Morlet wavelets for frequencies of interest (delta (1-4 Hz), alpha (8-14 Hz), low gamma (30-60 Hz), and high gamma (60-90 Hz). CSDs were then used to construct spatial filters via a dynamic imaging of coherent sources (DICS (Gross et al., 2001)) beamformer. This calculation returns a source power estimate over defined frequencies, from which the average absolute power was computed, giving one value per frequency range per source space vertex. Then, for each participant, the mean was computed over vertices within each ROI (see below), yielding an average power value per frequency range per ROI, allowing for statistical evaluation at the participant, group, and ROI level.

#### 3.2.1 Regions of Interest (ROIs)

##### Auditory Cortex (A1) and Secondary Auditory Cortex (A2)

With regard to auditory processing, we were most interested in Brodmann Areas BA41 and BA42. These correspond to the primary and secondary auditory cortices, respectively. Cortical parcellations were derived from the Destrieux cortical atlas (Destrieux et al., 2010). To define the boundaries of primary auditory cortices (A1), we combined two smaller subsections as provided in the Destrieux atlas, namely the anterior transverse temporal gyrus (Heschl’s Gyrus) and the transverse temporal sulcus. We defined secondary auditory cortices (A2) by combining the planum polare and planum temporale of the superior temporal gyrus, and the lateral aspect of superior temporal gyrus. Both A1 and A2 were analyzed with hemispheres combined (to measure overall auditory activation) as well as separated into left and right hemispheres (to assess the extent, if any, of oscillatory lateralization).

#### 3.2.2 Statistic evaluation of source space results

Effects of condition (*tinnitus, no tinnitus*) and hemisphere (*right, left*) on oscillatory power were investigated via a two-way analysis of variance (ANOVA) of a fitted ordinary least squares (OLS) linear model of group source space results. This analysis was run separately for each ROI. Post-hoc tests for multiple comparisons were performed using Tukey’s HSD (honestly significant difference) test.

All statistical analysis was performed using open-source software Python (version 3.6.1) and associated statistical packages and modules (e.g., scipy (Virtanen et al., 2020)).

## 4. Results

### 4.1 Audiometric Data

#### 4.1.1 Hearing Thresholds

Both earplug attenuation checks demonstrated that the hearing thresholds of the plugged ear were lower than those of the unplugged ear. After the plug was removed and participants had completed the Day 4 MEG measurement, another audiometric test was done to verify that hearing thresholds for both the plugged and unplugged ear recovered back to pre-plug levels (Figure 2). Paired-sample t-tests showed no statistical difference between participants’ hearing thresholds in the left or right ear before and after earplugging.

**Figure 1.**
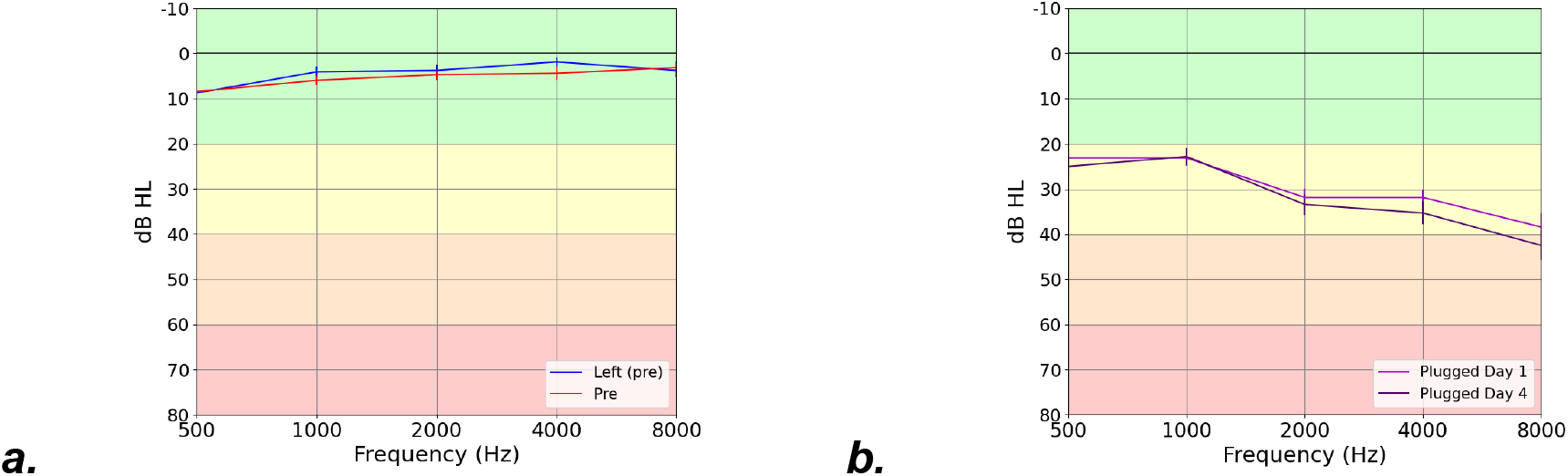
**(a)** Average baseline (i.e. pre-experiment) hearing thresholds for left (blue) and right (red) ears. **(b)** Average hearing thresholds for the plugged ear while the plug was inserted on day 1 (light purple) and day 4 (dark purple).

**Figure 2.**
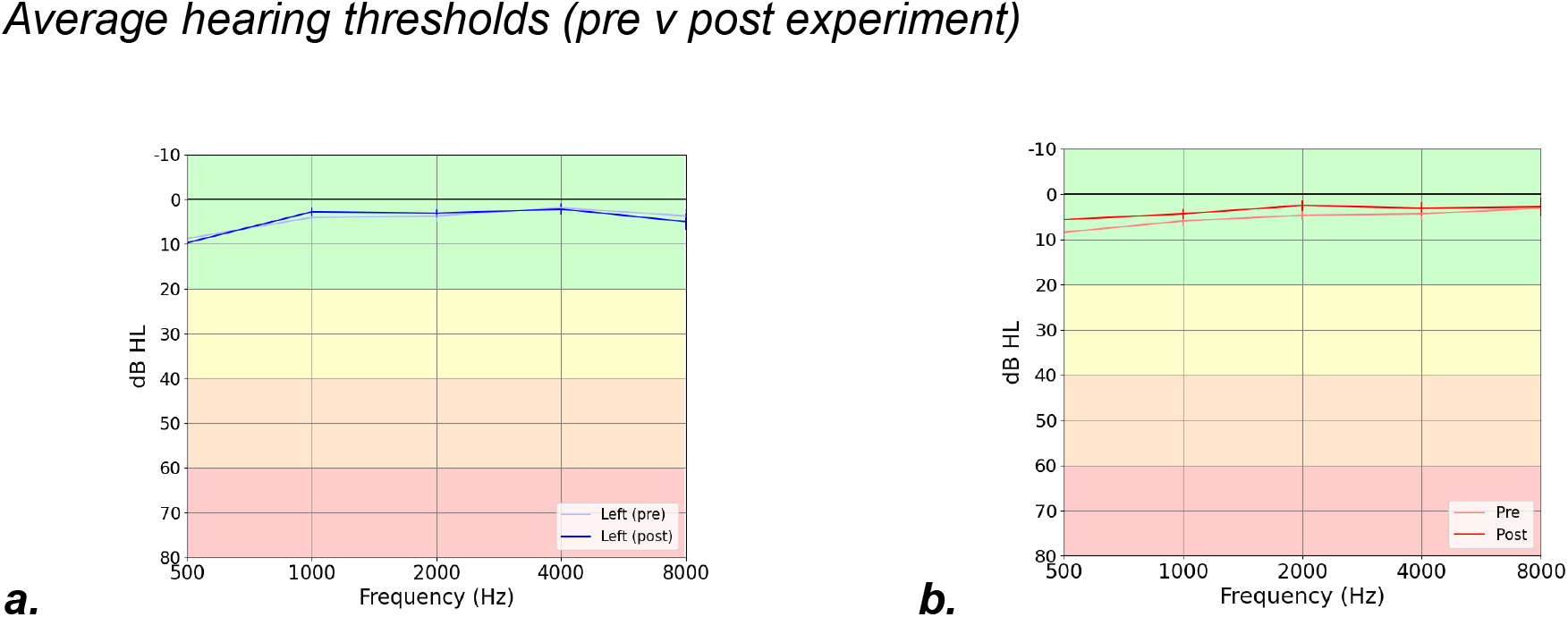
Average hearing thresholds for left **(a)** and right **(b)** ears. Paired-sample t-tests revealed no significant differences for the left or right when comparing pre- and post-experimental hearing thresholds (p > 0.1).

#### 4.1.2 Phantom Auditory Percepts

Of the 12 participants that completed the entire experiment, 11 participants developed unilateral phantom auditory percepts throughout the course of the four days, 10 experienced these percepts during the crucial Day 4 MEG measurement, and one participant did not experience any tinnitus-like noises at all. Most participants experienced pure tone phantom sounds which fell between 4000 and 8000 Hz, corresponding to the frequency range most attenuated by the specific model of silicone putty earplug used. Other characteristics of the phantom percepts are contained in Table 1. All 10 participants who were still experiencing phantom sounds during the tinnitus-condition resting state Day 4 measurement reported that these noises disappeared within seconds to minutes following earplug removal (on average, the sounds lingered for approximately 2 minutes, although it proved difficult for participants to confidently claim precisely when phantom sounds were completely attenuated).

#### 4.1.3 Hearing Loss vs Phantom Percept Frequency

We investigated the frequencies of participants’ subjective tinnitus-like percepts as a function of hearing loss (Figure 3). Pearson’s correlation revealed a weakly positive correlation between hearing loss and tinnitus percept frequency, although it remained statistically insignificant (r = 0.51, p = 0.11).

**Fig 3.**
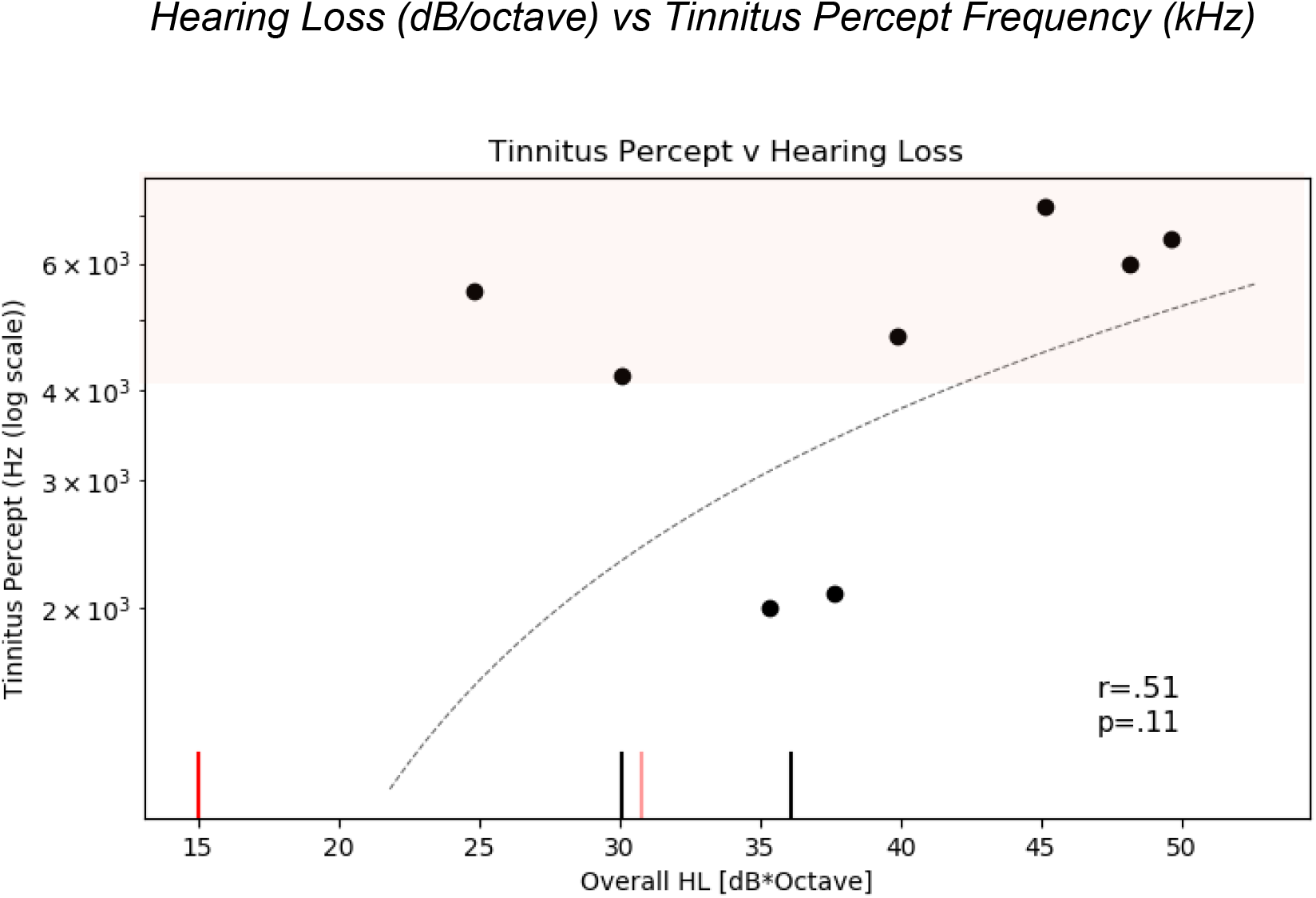
i. Subjective pitch judgment of tinnitus-like percept versus hearing loss. The red shaded area indicates the frequency range most attenuated by the earplugs (4-8 kHz). We therefore expected most tinnitus percepts to fall within this range. ii. The two vertical black lines on the x-axis denote two participants whose tinnitus-like percept took on more buzzing or noisy characteristics, unlike most participants who experienced pure tones or ringing. iii. The two vertical red lines represent participants who did not experience any phantom sounds during the tinnitus-condition measurement (*light red* = experienced pure tones during experiment but not during measurement, *bright red* = did not experience tinnitus).

### 4.2 Oscillatory Power Analysis

We investigated changes in cortical oscillatory power associated with the development of phantom auditory sounds after four days of unilateral deprivation. The average absolute power for delta (1-4 Hz), alpha (8-14 Hz), and gamma (30-60 Hz) bands were computed for each condition in primary (A1) and secondary auditory cortices (A2). For clarification, the **tinnitus** condition refers to MEG resting state measurements with ears plugged, no stimulus, and phantom sound perception, while the **silence** condition refers to resting state with earplugs, no stimulus, and no phantom sounds.

All participants who experienced phantom sounds reported that they were unilateral, and were only perceptible on the side of the plugged ear. We also compared contralateral and ipsilateral A1 and A2 activation during tinnitus and silence but found no correlation with the perceived side of the phantom sounds (p=0.15).

#### 4.2.1 Primary Auditory Cortex (A1)

##### Delta (1-4 Hz)

Two-way ANOVA revealed a significant effect of hemisphere on delta-band power (F(1,32)=4.141, *p*=0.04)), which also survived Tukey’s HSD test (*p*=0.04). Right primary auditory cortex showed greater activation levels in both the silence (*t*=3.269, *p*=0.013) and tinnitus conditions (*t*=2.807, *p*=0.023). No significant effect of condition was found.

##### Alpha (8-14 Hz)

The same analyses revealed no significant effect of condition or hemisphere on oscillatory power in the alpha frequency band. Additionally, no significant interaction was found between condition and hemisphere.

##### Gamma (30-60 Hz)

Two-way analysis of variance (ANOVA) revealed a significant effect of both condition (F(1,32)=6.535, p=0.016) and hemisphere (F(1,32)=5.491, p=0.025) on gamma-band cortical oscillatory power (Figure 4). The interaction between condition and hemisphere, however, was insignificant (F(1,32)=0.081, p=0.78). Post-hoc analysis of ANOVA results via Tukey’s HSD test for multiple comparisons revealed a significant increase of gamma power in the primary auditory cortices (A1) during the tinnitus condition (p=0.02) (Figure 4a). The effect of hemisphere on gamma power was observed, following correction for multiple comparisons (p=0.046). When comparing left and right A1 gamma activation levels via paired sample *t*-test, right primary auditory cortices were significantly higher in both the silence (*t*=3.384, *p*=0.01) and tinnitus conditions (*t*=2.504, *p*=0.04) (Figure 4b).

**Figure 4.**
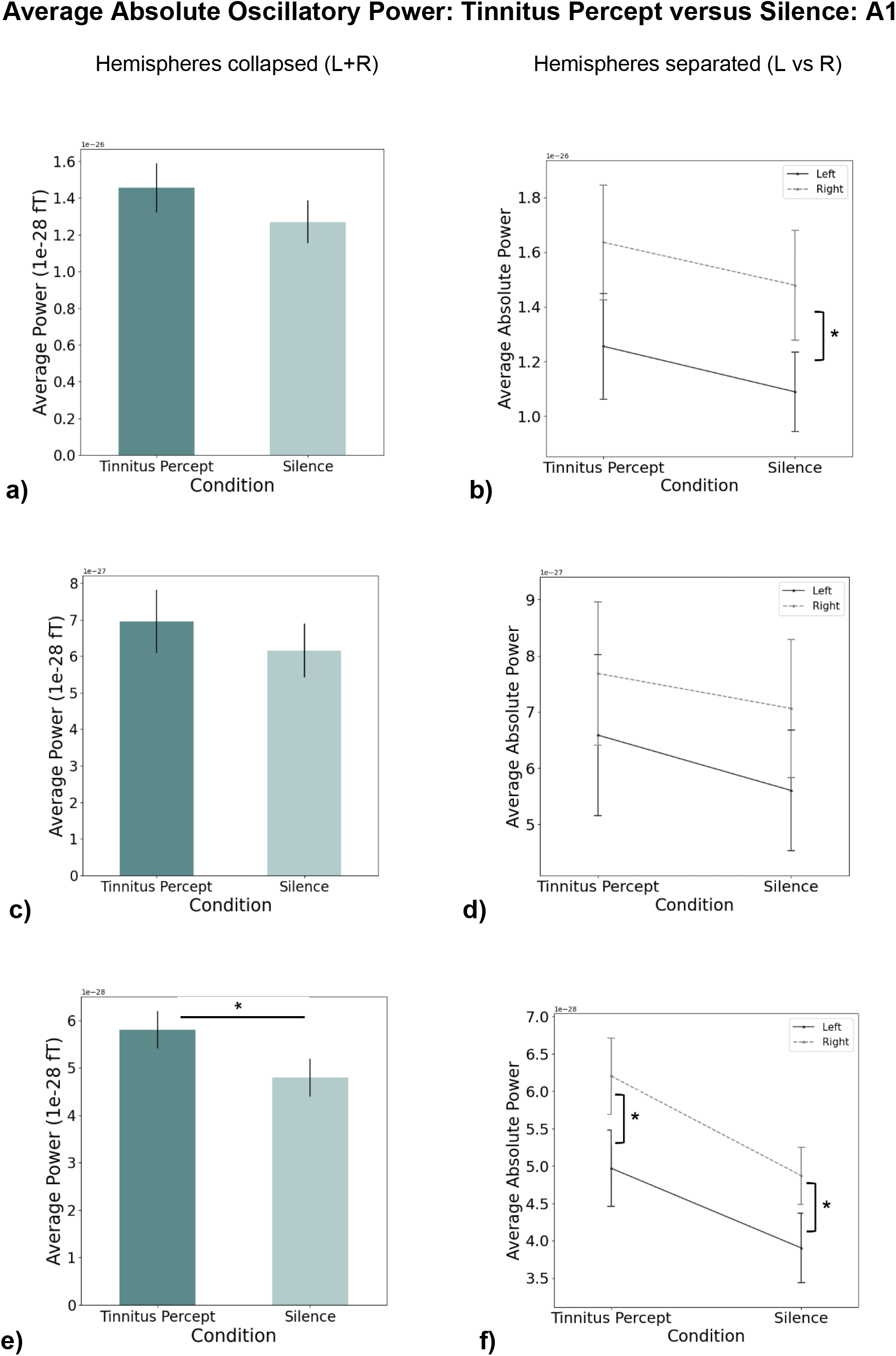
Power analyses results for delta (1-4 Hz) **(a**,**b)**, alpha (8-14 Hz) **(c**,**d)**, and gamma (30-60 Hz) **(e**,**f)** oscillatory activity in primary auditory cortex (A1). The left column shows combined (left and right) auditory cortex average absolute power 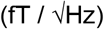, and the right column shows oscillatory power values separated between right and left auditory cortices. * *p<0.05*

## 5. Discussion

### 5.1 Increased auditory cortical synchrony following earplugging

Four days of unilateral deprivation was sufficient to elicit phantom auditory percepts in 10 of 11 participants (90.9%). Additionally, a significant increase in gamma power (30-60 Hz), localized to primary auditory cortex (A1 (left + right hemispheres)), was observed when participants heard phantom sounds compared to when they did not. This effect was observable when right and left auditory cortices were analyzed together, as well as when they were separated. No modulation of cortical delta or alpha was seen in A1 or A2 as a result of unilateral deprivation. Interestingly, group-averaged hemispheric data revealed significantly higher delta and gamma power in right primary auditory cortex compared to left, both during phantom auditory perception as well as during silence, regardless of the earplugging side.

### 5.2 Mechanistic role of gamma oscillations

Group-averaged A1 gamma power was significantly higher in the tinnitus condition when compared with the silent condition. This result is in line with increased A1 gamma activity observed with MEG in both chronic tinnitus perception (Weisz et al., 2005; Van Der Loo et al., 2009) and acute tinnitus perception (Ortmann et al., 2010; Lan et al., 2020). Apart from tinnitus, increased auditory gamma power is thought to be a neural correlate of acoustic stimuli processing in healthy populations, after evidence from MEG (Pantev et al., 1991; Palva et al., 2002) and EEG (Galambos et al., 1981; Bertrand, O. & Pantev, C., 1994) observed gamma-band synchrony in auditory cortices. However, it remains to be resolved whether the gamma-band generators involved in acoustic perception, chronic tinnitus, and acute tinnitus are the same. It is likely that distinct neural networks modulate auditory gamma-band synchrony in these three cases, given that altered cortico-cortical connectivity patterns are found in chronic tinnitus populations compared with healthy controls (Cacace, A.T., 2003; Madoux et al., 2012), as well as the inherent difference between exogenous and endogenous sounds. In some chronic tinnitus neuroimaging studies, increased A1 gamma power correlates with increased delta (1-4 Hz) and decreased alpha (8-14 Hz) (Weisz et al., 2007). This pattern has not yet been observed in unilateral acute tinnitus populations, including the present study. The oscillatory changes seen in chronic tinnitus groups may be correlated to phantom percept chronification (as opposed to “mere” phantom perception), although more experiments monitoring neural oscillations associated with tinnitus onset and development would be necessary to confidently establish this relation.

In our study, the observed upregulation of neural activity in response to a decrease in external sensory input correlated with subjective reports of phantom sound perception (i.e., increased *internal* sensory input). This finding could, in principle, be explained by thalamocortical dysrhythmia, a theory which suggests that increased gamma power in the auditory cortices of tinnitus sufferers is the result of an imbalance at the level of the thalamus due to reduced cochlear input, which ultimately results in an “edge effect” at borders of auditory cortical tonotopic organization representing the affected frequency ranges, disinhibiting normally silent neurons (Llinas et al., 1999, 2005). Homeostatic plasticity, another current theory in the neuroscience of tinnitus, argues that a reduction in cochlear input leads to an increase in central auditory gain as a way to recompensate mean neural activity levels to their original values (i.e., before loss of cochlear input) (Yang et al., 2011; Schaette and Kempter, 2006, 2009, 2012; Brotherton et al., 2019). While the present study cannot definitively point to thalamocortical dysrhythmia or homeostatic plasticity as the theoretical mechanism behind our results, it does indeed demonstrate that four days of earplug use is enough to engage whatever that mechanism might be.

### 5.3 Delta and Gamma-Band Lateralization

Gamma power in primary auditory cortex was not only significantly higher in the tinnitus condition versus the silent condition but was also significantly higher in the right hemisphere versus left, regardless of condition. Similarly, delta waves were also stronger in the right hemisphere, during both conditions. Lateralization of certain auditory functions in healthy humans is well established, with the left hemisphere differentially contributing to language processing (Devlin et al., 2003) and the right hemisphere involved more in musical or tonal processing (Zatorre et al., 2002). Additionally, auditory cortices respond preferentially to the contralateral ear’s input (Pantev et al., 1998). With regards to unilateral tinnitus patients, M/EEG have shown positive correlations between contralateral A1 gamma activation and unilateral tinnitus intensity (Weisz et al., 2005; Van der Loo et al., 2009). However, in a study using fluoro-deoxyglucose positron emission tomography (FDG-PET), Geven et al. demonstrated no tinnitus-specific lateralization of A1 or A2 activity, although both the tinnitus groups (left and right unilateral percept) and control group exhibited left hemispheric dominance in A1 and right hemispheric dominance in A2 (Geven et al., 2014). Given that A1 and A2 are just one level of a deeper, more complicated system, future tinnitus studies may benefit from recording neuronal activity from various sites along the auditory pathway simultaneously, which would place cortical oscillations in a fuller context of central auditory functioning.

### 5.4 Role of Attention in Tinnitus

Attention is a contributing factor which should not be underestimated when considering not only the development of tinnitus, but also its chronification and potential negative impact on sufferers’ well-being. One theory accounts for the relation between tinnitus chronification and attention by suggesting that a predictive coding mechanism, present in people with healthy hearing (Näätänen et al., 1978, 2007; Gagnepain et al., 2012), acts to upregulate auditory attention when the brain detects a discrepancy between what is expected and what is perceived (Robert et al., 2013). During tinnitus perception, however, this attention may become entrenched, leading to chronification. Other research suggests altered functional connectivity patterns between auditory and non-auditory areas, including frontal and parietal areas which are implicated in executive and attentional control, which may pivot auditory attention toward tinnitus perception, cyclically exacerbating the experience (Haab et al., 2009; Trevis et al., 2016). Given that the participants who completed this study only heard phantom sounds for at most four days, auditory attention entrenchment was unlikely.

Unlike the participants included in the analysis of this study, one participant (P13) did report that their tinnitus-like percepts failed to disappear within minutes after earplug removal. While this was not unexpected, a follow-up meeting with P13 one week after earplug removal revealed that the phantom sounds persisted. P13 was referred to otolaryngological experts at FAU’s ENT clinic, who ruled out the presence of accumulated earwax or other physical obstructions which could continue to attenuate incoming noise. P13’s hearing thresholds were measured again, from 125 Hz to 16 kHz, with both ears demonstrating acoustic detection statistically indistinguishable from pre-earplugging levels. From an audiological perspective, no abnormalities were found within P13’s checkups. That being said, P13 was also 26 years old at the time of the experiment, so hidden hearing loss should not necessarily be ruled out. Two weeks after plug removal, authors 1, 3, and 5 met with P13 to understand their experience, both physical and emotional. According to P13 at this time, the same high-frequency (∼ 7 kHz) tonal phantom sounds that they heard during the experiment were still present, particularly at night or in moments of decreased ambient decibel levels. P13 also reported negative feelings of frustration or annoyance associated with the percept, although these did not hinder their professional or personal lives. Attentional strategies were explained and suggested (i.e., shift one’s focus away from the auditory percept) to P13. Due to P13’s report of persistent percepts, the study was not continued, and therefore our sample size remained much smaller than anticipated.

When investigating and inducing tinnitus-like experiences in a laboratory setting, how attention is guided over the course of an experiment may substantially influence behavioral outcomes. 90.9% of our participant pool (11/12) experienced tinnitus during the experiment, which is quite high when comparing the behavioral outcomes of this study with previous unilateral deprivation experiments. For example, when pooling participants from two unilateral deprivation studies, Brotherton et al. found 68% of participants heard phantom sounds in a previous study, despite participants undergoing a deprivation period of almost twice as long (one week versus four days) (Schaette et al., 2012; Brotherton et al, 2019). The main difference between these studies and the present study lies in their experimental approaches, namely that Schaette et al. never mentioned the word *tinnitus* to the participants during the experiment or its description, intentionally attempting to avoid whatever negative connotations participants could associate with the term. In the present study however, not only was the term *tinnitus* included in the name of the experiment (as described in participant recruitment calls and consent forms), but participants were explicitly asked to make a note of phantom sounds and their qualities (e.g., loudness, timbre, pitch, time of onset, etc.). It is likely that this sustained auditory attention over the course of the experiment contributes to behavioral outcome discrepancies between this study and other unilateral deprivation studies. This role of attention in the perception, maintenance, and severity of tinnitus may, however, prove to be a crucial variable in tinnitus management, the modulation of which could be leveraged so as to attenuate associated loudness and distress.

### 5.5 Limitations

In this study, the reduction in cochlear input induced via earplug use can be considered an *artificial* hearing loss, and therefore one should be careful not to assume that the changes along the auditory pathway observed in this study precisely parallel neural mechanisms underlying true hearing loss (e.g., noise trauma, cell degeneration, etc.).A limitation common to deprivation studies is in verifying that participants do indeed follow experimenters’ instructions and wear the earplug(s) for the required time. Strategies to resolve this potential confound in the present study were considered, but ultimately no practical solution was found. This was one reason we encouraged participants to keep a daily journal, as a method to “monitor” their genuine participation. However, we are confident that the current protocol was successful, given that all but one participant reported the development of tinnitus over the course of the experiment.

A further limitation of this study was that analyses were confined to primary and secondary auditory cortices only, corresponding to our initial hypotheses. Future connectivity analyses will aim to reveal communication between auditory and non-auditory cortical areas. A follow-up experiment could also benefit from novel techniques which allow for the investigation of deeper brain structures.

Finally, another limitation to this study was the difficulty in obtaining a larger sample size of participants. We were only able to recruit and complete the experiment with about a quarter of our original target sample size of 60 participants. This was due mostly to P13’s complications as mentioned above, which necessitated a redrafting of the ethics approval. Unfortunately, by this time, further recruitment and experimental data collection was hampered by both logistical challenges presented by the pandemic, as well as demolition carried out on the building which housed our MEG.

## 6. Conclusion

To the best of our knowledge, this study is the first MEG experiment in combination with auditory deprivation via earplug use. By wearing an earplug in one ear continuously for four days, we simulated an artificial moderate hearing loss in 12 healthy hearing participants. During the deprivation period, all but one participant experienced tinnitus-like phantom auditory sensations. Using MEG, we found increased gamma oscillations source-localized to primary auditory cortex when participants were experiencing these percepts compared to when they were not, while modulations of delta and alpha oscillations were not detected. It is reasonable to suggest that the observed changes in oscillatory activity in this study reflect increased neural synchrony as a result of reduced unilateral cochlear input. The rapidity with which these deprivation-induced plastic changes of auditory processing take place not only demonstrates the utility of such an experimental paradigm, but sheds light on how quickly adaptive auditory plasticity is engaged. A better understanding of auditory plasticity in general, and which cerebral mechanisms lead to phantom auditory percepts in particular, will be fundamental to future research and therapy designed to aid those suffering from tinnitus.

