## Supplemental Figure 1 for "Cortical Activity Associated With Acute Development of Phantom Auditory Percepts via Unilateral Deprivation"

### Supplementary Material

#### Average Absolute Oscillatory Power: Tinnitus Percept versus Silence: A2

Hemispheres collapsed (L+R) Hemispheres separated (L vs R)

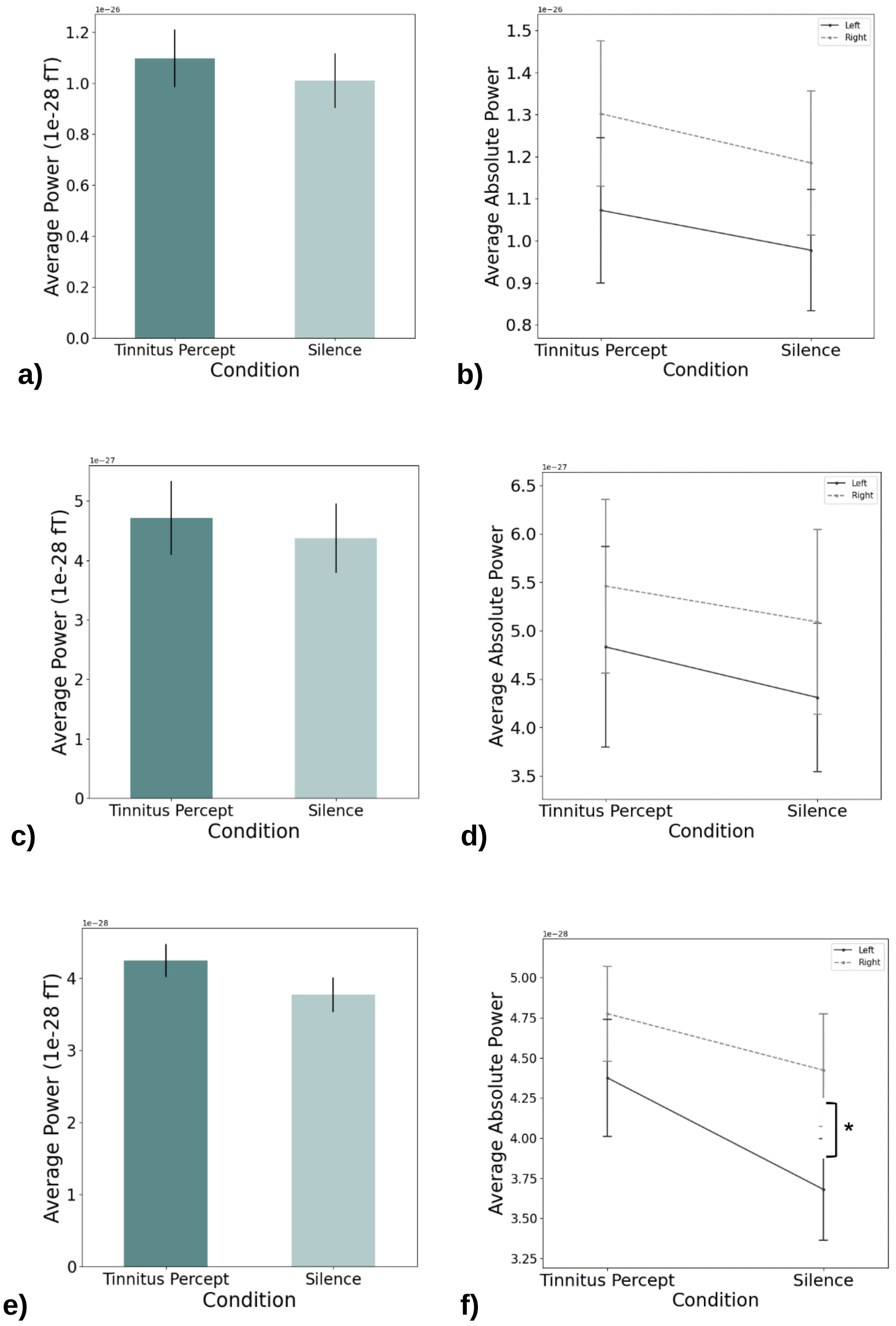


**Figure S1.** Power analyses results for delta (1-4 Hz) **(a,b)**, alpha (8-14 Hz) **(c,d)**, and gamma (30-60 Hz) oscillatory activity in secondary auditory cortex (A2). The left column shows combined (left and right) auditory cortex average absolute power (fT / √Hz) (tinnitus = dark green; silence = light green. The right column shows oscillatory power values separated between right A2 (dotted line) and left A2 (solid line). A paired-sample t-test confirmed significantly higher gamma power during silence in right A2 than left A2. This activation asymmetry was not observed during the tinnitus condition. * $p<0.05$
